# Local and sex-specific biases in crossover vs. noncrossover outcomes at meiotic recombination hotspots in mouse

**DOI:** 10.1101/022830

**Authors:** Esther de Boer, Maria Jasin, Scott Keeney

## Abstract

Meiotic recombination initiated by programmed double-strand breaks (DSBs) yields two types of interhomolog recombination products, crossovers and noncrossovers, but what determines whether a DSB will yield a crossover or noncrossover is not understood. In this study we analyze the influence of sex and chromosomal location on mammalian recombination outcomes by constructing fine-scale recombination maps in both males and females at two mouse hotspots located in different regions of the same chromosome. These include the most comprehensive maps of recombination hotspots in oocytes to date. One hotspot, located centrally on chromosome 1, behaved similarly in male and female meiosis: crossovers and noncrossovers formed at comparable levels and ratios in both sexes. In contrast, at a distal hotspot crossovers were recovered only in males even though noncrossovers were obtained at similar frequencies in both sexes. These findings reveal an example of extreme sex-specific bias in recombination outcome. We further find that estimates of relative DSB levels are surprisingly poor predictors of relative crossover frequencies between hotspots in males. Our results demonstrate that the outcome of mammalian meiotic recombination can be biased, that this bias can vary depending on location and cellular context, and that DSB frequency is not the only determinant of crossover frequency.

## INTRODUCTION

During meiosis, homologous chromosomes (homologs) pair and undergo reciprocal DNA exchanges (crossovers), which are required for proper chromosome segregation and which promote genetic diversity. In many eukaryotes, including mammals, both homolog recognition and crossover formation involve homologous recombination initiated by SPO11-generated DNA double-strand breaks (DSBs) (Lam and Keeney 2014). Recombination at multiple chromosomal positions supports homolog pairing but only a subset of DSBs become interhomolog crossovers; the rest give rise to interhomolog noncrossovers or genetically silent sister-chromatid recombination (Schwacha and Kleckner 1994; Allers and Lichten 2001; Hunter and Kleckner 2001; Goldfarb and Lichten 2010). Studies in the budding yeast *Saccharomyces cerevisiae* indicate that the two types of interhomolog recombination products arise through distinct pathways (reviewed in Youds and Boulton 2011). Most noncrossovers appear not to involve Holliday junction intermediates, but instead are thought to arise through synthesis-dependent strand annealing. In contrast, most crossovers are formed by resolution of double Holliday junction intermediates through the canonical DSB repair pathway and require a group of proteins collectively known as the ZMM proteins (Zip1, Zip2, Zip3, Mer3 and Msh4/Msh5) as well as the MutL-homologs Mlh1 and Mlh3 for their formation. A subset of crossovers is formed by an alternative pathway(s) involving Holliday junction resolution by distinct resolvases including Mus81-Mms4 or Yen1. Studies in other organisms indicate that the existence of distinct pathways for forming crossovers vs. noncrossovers is a conserved feature of meiosis (e.g., (Cole et al. 2014)), although detailed operation of these pathways often differs between taxa (Youds and Boulton 2011).

Recombination is regulated to ensure that each chromosome receives at least one crossover and that multiple crossovers on the same chromosome (if they occur) tend to be widely and evenly spaced (Jones and Franklin 2006). Studies in many species have focused on how multiple recombination events on individual chromosomes are regulated with respect to one another (e.g., de Boer et al. 2006; Libuda et al. 2013), but less attention has been paid to the question of whether recombination plays out similarly at all genomic locations. Cytological data in mouse suggest that there are significant differences between large (tens of megabases) chromosomal domains in the likelihood that a DSB will give rise to a crossover versus noncrossover outcome (de Boer et al. 2006), but direct molecular tests of these differences are lacking.

Most crossovers in humans and mice occur in narrow (∼1–2 kb wide) regions termed hotspots, which overlap preferred sites of SPO11 DSBs (Brick et al. 2012; Baudat et al. 2013). Individual crossover hotspots have been extensively characterized by analysis of human and mouse sperm DNA (Jeffreys et al. 2001; Guillon and de Massy 2002; Bois 2007; Cole et al. 2010). Noncrossovers are less well characterized, but where examined occur in the same hotspots where crossovers occur (Jeffreys and May 2004; Guillon et al. 2005; Cole et al. 2010; Sarbajna et al. 2012). While crossovers are readily identified because they exchange large DNA segments, noncrossovers are only detectable if a sequence polymorphism(s) is copied from the intact homolog during DSB repair. Because noncrossover gene conversion tracts are short, detection is highly dependent on the location of polymorphisms relative to DSBs, which differs between hotspots (Guillon et al. 2005; Cole et al. 2010). Not surprisingly then, the ratios of detectable noncrossovers to crossovers vary widely between hotspots, ranging from more than a 10-fold excess of crossovers to a 10-fold excess of noncrossovers, the latter of which is close to the genome average estimated by cytology of DSB and crossover markers (Holloway et al. 2006; Baudat and de Massy 2007b; Cole et al. 2010; Sarbajna et al. 2012). Thus, to what degree the variation in crossover:noncrossover ratios is a consequence of technical limitations or instead reflects genuine differences in recombination outcome has usually been unclear. Moreover, a recent study of DSB formation in human males emphasized recombination outcome similarities between genomic locations by focusing on DSB frequency as the principal determinant of crossover frequency (Pratto et al. 2014).

Comparatively little is known about recombination in females. In mouse pedigree studies of recombination on chromosomes 1 and 11, sex-specific crossover hotspots were identified and some shared hotspots showed male versus female differences in crossover frequencies (Paigen et al. 2008; Billings et al. 2010). Because pedigree analysis only detects crossovers, it is uncertain whether this sex-specific variation in recombination results from differences in DSB formation, recombination outcome, or both. Molecular studies of recombination in females have been limited, given the technical challenges of studying recombination in the fetal ovary where meiosis occurs. Further, oocytes are embedded within somatic tissue rather than being present in a ductal lumen as for sperm, and the number of oocytes recovered from a female is much less than the number of sperm from a male. Thus far, only one mouse hotspot (*Psmb9*, initially identified as a female-specific hotspot (Shiroishi et al. 1990)) has been directly assayed for crossovers and noncrossovers in both males and females, but available data in females are limited to relatively few recombinant molecules and/or assays of noncrossover gene conversion at just one or a small number of polymorphisms (Guillon et al. 2005; Baudat and de Massy 2007a; Cole et al. 2014).

In this study, we generated comprehensive, high-resolution maps of crossovers and noncrossovers in both males and females at two previously uncharacterized hotspots located in distinct regions on the same chromosome. Examining the same hotspots in both sexes allowed us to analyze sex-specific variation in recombination outcomes free of confounding effects attributable to the positions of scoreable sequence polymorphisms. We also compared relative crossover frequencies with published estimates of relative DSB frequencies. The findings reveal striking sex-specific and local differences between hotspots in the likelihood that a DSB will give rise to a crossover vs. noncrossover recombination product.

## RESULTS

### Hotspot selection

To test the hypothesis that there can be regional and sex-specific variation in recombination outcomes, we selected hotspots located in regions displaying different propensities towards crossover formation. The central approximately one-third of mouse chromosome 1 shows a similar crossover preference in males and females, based on relative frequencies of MSH4 foci (a cytological marker of early recombination intermediates) and MLH1 foci (a marker of crossovers) (de Boer et al. 2006). In contrast, males generate crossovers preferentially in centromere-distal subtelomeric regions (Shifman et al. 2006; Paigen et al. 2008). This is not matched by a higher frequency of MSH4 foci, suggesting that male predilection toward subtelomeric crossovers reflects a regional bias in the crossover:noncrossover ratio, not simply more DSBs (de Boer et al. 2006). If so, this bias is at least partially sex-specific (de Boer et al. 2006; Shifman et al. 2006; Paigen et al. 2008).

We identified a hotspot from the central region of chromosome 1 using recombinant inbred (RI) strains. These strains contain patchworks of different genomic segments from two parental inbred strains, boundaries of which are determined during the generations of inbreeding by crossovers at potential hotspots (Bois 2007) (**Figure S1**). Sixteen candidate hotspots were initially examined, of which one proved appropriate for further analysis because it has a suitable polymorphism density and lacks repeats that would compromise PCR amplification (see **Supplemental Information**). This central hotspot, located at 78.59 Mbp on chromosome 1 (**Figure 1A**), has a polymorphism density of 0.97% between the C57BL/6J (hereafter B6) and A/J haplotypes (30 polymorphisms across 3094 bp; **Table S1**). A second hotspot, at 185.27 Mbp on chromosome 1 (**Figure 1A**, “distal hotspot”), was selected from a whole-genome map of recombination initiation sites identified by deep sequencing of ssDNA bound by the RAD51 or DMC1 strand exchange proteins (Smagulova et al. 2011). The polymorphism density between B6 and A/J haplotypes at this hotspot is 0.89% (36 polymorphisms across 4048 bp; **Table S2**).

**Figure 1.**
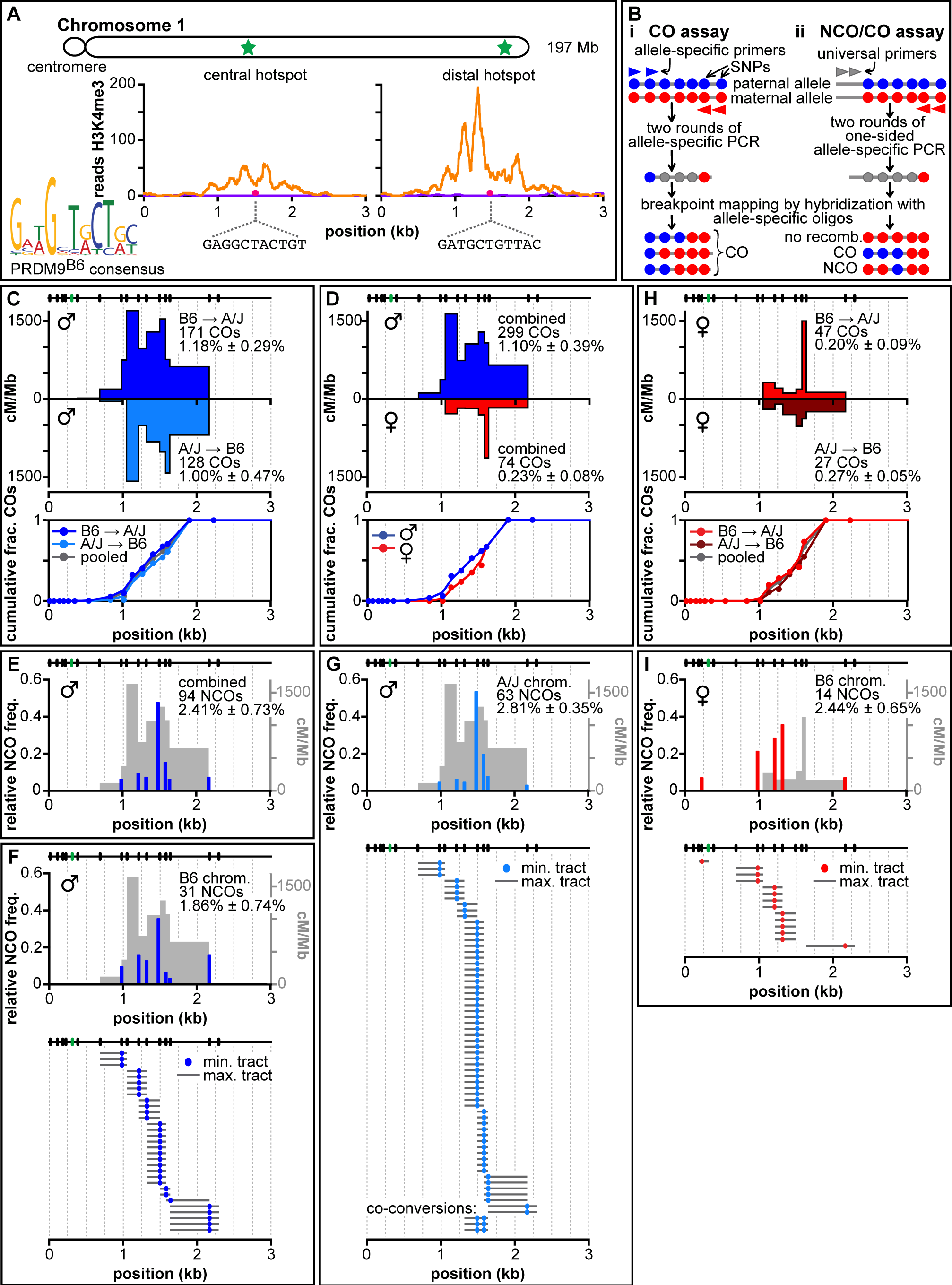
Recombination at the central hotspot in males and females. **A.** Overview of recombination hotspots in this study. Top: Schematic of mouse chromosome 1 with hotspot positions indicated as green stars. The central hotspot is located between bp 78,589,305-78,592,399. The distal hotspot is located between bp 185,265,469-185,269,517 (build 38 for both). Bottom: Both hotspots show H3 lysine 4 trimethyl (H3K4me3) ChIP-seq signals (Baker et al. 2014) in testes of mice expressing the B6 version of PRDM9 (orange trace) but not the PRDM9 version found in the CAST/EiJ strain (purple trace, which mostly overlaps the horizontal axis). The 11-bp motif predicted to bind B6 (and A/J)-encoded PRDM9 (bottom left) is found in both hotspots (pink dots) and is identical in the B6 and A/J strains for each hotspot. Depicted regions in these and all subsequent hotspot graphs: 78,589,447–78,592,447 for the central hotspot; 185,265,656–185,268,656 for the distal hotspot. **B.** Assays to amplify and identify crossovers and noncrossovers. Filled circles represent sequence polymorphisms (red or blue for the two parental genotypes, gray for amplified but not-yet-defined internal polymorphisms); arrowheads represent PCR primers (red or blue for allele-specific primers, gray for universal). (i) In the crossover (CO) assay, two sequential rounds of allele-specific PCR selectively amplify recombinant DNA molecules from small pools of sperm or oocyte DNA of an F1 hybrid animal. Recombination frequencies are estimated from the observed fraction of pools that yield amplification products. Next, internal polymorphisms in each amplified recombinant DNA molecule are genotyped by hybridization with ASOs to map the location of the crossover breakpoint. (ii) In the noncrossover/crossover (NCO/CO) assay, smaller pools of sperm or oocyte DNA are amplified using nested primers that are specific for one haplotype in combination with nested universal primers. In contrast to the crossover assay, both non-recombinant and recombinant DNA molecules are amplified, with the majority (>95%) being non-recombinants. Subsequent hybridization of amplification products to ASOs that are specific for alleles from the non-selected haplotype identifies pools containing crossovers and noncrossovers. **C.** Crossovers in males. Crossover breakpoint maps (top graph) are shown for crossover molecules amplified with allele-specific primers in the B6 to A/J orientation (dark blue) or the A/J to B6 orientation (light blue). Cumulative crossover distributions are shown below. Tested polymorphisms are indicated as ticks at the top. Multiple polymorphisms contained within a single ASO are indicated as a single green tick. Numbers of observed crossovers and Poisson-corrected crossover frequencies (± SD) are indicated. **D.** Similar distributions of crossover breakpoints in males and females. Data from both orientations of the allele-specific PCR were pooled separately for males (blue) and females (red). **E.** Total noncrossovers in males. Total relative noncrossover frequencies from all four orientations of the PCRs (normalized for co-conversions) at the tested polymorphisms are shown as blue bars. The crossover breakpoint map in males is shown for comparison (light gray). Number of total observed noncrossovers and Poisson-corrected total noncrossover frequency (± SD) are indicated. **F, G.** Noncrossovers on the B6 (F) and A/J (G) chromosome in males. Top: relative noncrossover frequencies (normalized for co-conversions) on the B6 (F) and A/J (G) chromosome at the tested polymorphisms are shown as blue bars. The crossover breakpoint map in males is shown for comparison (light gray). Number of observed noncrossovers and Poisson-corrected noncrossover frequency (± SD) are indicated. Bottom: noncrossover gene conversion tracts on the B6 (F) and A/J (G) chromosomes. **H.** Crossovers in females. Crossover breakpoint maps (top graph) and cumulative crossover distributions (bottom graph) are shown for the B6 to A/J orientation (light red) and the A/J to B6 orientation (dark red) of allele-specific PCR. **I.** Noncrossovers in females. Relative noncrossover frequencies and noncrossover gene conversion tracts from PCR in the universal-to-B6 orientation are presented as for males.

A principal determinant of DSB hotspot location is the meiosis-specific PRDM9 protein, which contains a histone methyltransferase domain and a DNA binding domain comprising an array of Zn-finger modules whose DNA binding specificity evolves rapidly (Baudat et al. 2010; Myers et al. 2010; Grey et al. 2011; Brick et al. 2012). Both the central and distal hotspots contain a match in both the B6 and A/J strains to the motif predicted to bind the B6-encoded PRDM9 (A/J has the same *Prdm9* allele), and PRDM9-dependent histone H3 lysine 4 trimethylation is present at both hotspots as determined by chromatin immunoprecipitation (Baker et al. 2014) (**Figure 1A**). Thus, both hotspots are predicted to be active sites for DSB formation in B6 x A/J F1 hybrids.

### Recombination outcomes at the central hotspot in males

#### Crossovers

We examined crossing over in the central hotspot in males by allele-specific PCR on sperm DNA from B6 x A/J F1 hybrids (**Figure 1Bi**). Crossover molecules were selectively amplified from pools of sperm DNA by two rounds of PCR with nested forward primers specific for one parental haplotype, combined with nested reverse primers specific for the other parental haplotype (Jeffreys and May 2003). Crossover breakpoints in the amplified recombinant molecules were subsequently mapped by allele-specific hybridization at 18 polymorphisms across the hotspot. We retrieved 299 crossover molecules from a total input of ∼32,000 haploid genome equivalents, for a Poisson-corrected overall frequency of 1.10% per haploid genome (**Table 1**). No crossovers were detected in somatic DNA controls (frequency < 0.004%). Crossovers showed similar frequencies and breakpoint distributions for both orientations of allele-specific primers (i.e., B6 forward primers plus A/J reverse, or the converse; **Figure 1C**). The similar breakpoint distributions imply that recombination initiates at similar frequency on both haplotypes in this F1 hybrid strain, as inferred previously at other hotspots (Jeffreys and Neumann 2002; Cole et al. 2010). Combining data from both primer orientations, crossover activity spanned 1.8 kb, with an average of 627 cM/Mb and peak of 1638 cM/Mb (♂, **Figure 1D).** These values place this hotspot among the most active hotspots for crossovers characterized in male mice, comparable to *Psmb9* (1.1% crossover frequency, 1300 cM/Mb peak activity (Guillon and de Massy 2002)).

#### Noncrossovers

To detect noncrossovers, we used nested PCRs that were allele-specific on just one side, i.e., with primers for one of the parental haplotypes opposed to “universal” primers that amplify both haplotypes (**Figure 1Bii**) (Jeffreys and May 2004). Recombinant DNA molecules amplified from small pools of sperm DNA were detected by hybridization to allele-specific oligonucleotides directed against the non-selected parental haplotype. From all four possible orientations of allele-specific primers, we recovered 94 noncrossovers from a total input of ∼4200 haploid genome equivalents, for a Poisson-corrected overall frequency of 2.4% per haploid genome (**Figure 1E**, **Table 1, Table S1**). Noncrossovers occurred across the same region as crossovers and the peak of noncrossover activity overlapped the center of the crossover distribution (**Figure 1E-G**). The polymorphism showing the highest noncrossover frequency was adjacent to the PRDM9 motif (relative noncrossover frequencies of 0.36 on the B6 chromatid and 0.54 on the A/J chromatid). The minimum gene conversion tract of each noncrossover was measured by considering only the polymorphism(s) involved plus the segments in between if more than one polymorphism was converted; the maximum conversion tract includes the distance to the nearest flanking polymorphisms that were not converted (**Figure 1F and 1G**). For the combined noncrossovers on both the B6 and A/J chromosomes the average minimum and maximum conversion tracts were 4 and 304 bp, respectively. Both the distribution of noncrossovers across the hotspot as well as the conversion tract lengths are similar to reports for other hotspots (Baudat and de Massy 2007b; Cole et al. 2010). Co-conversions were rare: only three noncrossovers converted more than one polymorphism, all involving the same two polymorphisms near the hotspot center, with minimum and maximum tract lengths of 79 and 315 bp, respectively (**Figure 1G**). These co-conversions were independent events as they arose from separate sperm pools.

Importantly, this assay also detects crossovers (**Figure 1Bii**). We recovered 47 crossover molecules for a Poisson-corrected overall frequency of 1.15% (**Table 1**). Agreement of this value with that from the crossover-specific assay validates direct quantitative comparison between them. Each noncrossover recombination event generates a single recombinant DNA molecule from four chromatids, whereas each crossover event generates two (reciprocal) recombinants (Cole et al. 2014). Thus, the relative numbers of detectable recombinants in the noncrossover/crossover assay translate to a per-meiosis ratio of 1 crossover to 4.2 noncrossovers (crossovers: 2.30% of meioses (2 × 1.15%); noncrossovers: 9.64% of meioses (4 × 2.41%)). This is lower than the ∼1:10 ratio estimated for genome average and observed at the well-characterized *A3* hotspot (Baudat and de Massy 2007b; Cole et al. 2010).

### Recombination outcomes at the distal hotspot in males

#### Crossovers

In the crossover-specific assay, we recovered 269 crossover molecules at the distal hotspot from ∼93,000 haploid genome equivalents of sperm DNA from B6 × A/J F1 hybrids, for a Poisson-corrected overall frequency of 0.36% (**Figure 2A and 2B**; **Table 1**). Crossover breakpoints showed similar distributions for both PCR orientations (**Figure 2A**), again implying equivalent DSB frequencies on both haplotypes. No crossovers were detected in somatic DNA controls (frequency < 0.004%). Breakpoint mapping using hybridization to allele-specific oligonucleotides (ASOs) at 25 polymorphisms across the hotspot indicated that crossovers occurred across a 2.7 kb region, with most breakpoints in the central 1 kb. The hotspot averaged 116 cM/Mb, peaking at 361 cM/Mb (♂, **Figure 2B**). Although not as active for crossing over as *Psmb9* or the central hotspot, the distal hotspot is very active in males and is similar to *A3* (0.26% crossover frequency in B6 x DBA F1 hybrids) (Guillon and de Massy 2002; Cole et al. 2010).

**Table 1.** Summary of recombination outcomes at the central and distal hotspots

|  | ♂ |  | ♀ |  |
| --- | --- | --- | --- | --- |
|  | Central | Distal | Central | Distal |
| Crossover frequency from CO assay<sup>1</sup> (no./total)<sup>2</sup> | 1.10% ± 0.39%<br>(299/32,290) | 0.36% ± 0.03%<br>(269/93,102) | 0.23% ± 0.08%<br>(74/29,438) | <0.007%<br>(0/15,013) |
| Crossover frequency from NCO/CO assay<sup>1</sup> (no./total)<sup>2</sup> | 1.15% ± 0.31%<br>(47/4177) | 0.33% ± 0.12%<br>(25/8007) | 0.34% ± 0.24%<br>(2/584) | <0.017%<br>(0/5994) |
| Noncrossover frequency from NCO/CO assay<sup>1</sup> (no./total)<sup>2</sup> | 2.41% ± 0.73%<br>(94/4177) | 1.53% ± 0.44%<br>(115/8007) | 2.44% ± 0.65%<br>(14/584) | 2.26% ± 0.80%<br>(122/5994) |
| Crossover:noncrossover<sup>3</sup> | 1:4.2 | 1:9.3 | 1:14.4<sup>4</sup> | <1:266 |
| SSDS reads<sup>5</sup> | 2819 | 9247 | NA<sup>6</sup> | NA<sup>6</sup> |

<sup>1</sup>Per haploid genome, Poisson-corrected, ± SD.
<sup>2</sup>Observed number of recombinants and total haploid genome equivalents analyzed.
<sup>3</sup>Ratio calculated from per-meiosis recombination frequencies [(noncrossover x 4) ÷ (crossover x 2)] from the NCO/CO assay.
<sup>4</sup>Note that this ratio is based on a small number of events.
<sup>5</sup>From Brick et al. (2012).
<sup>6</sup>NA, data not available.

**Figure 2.**
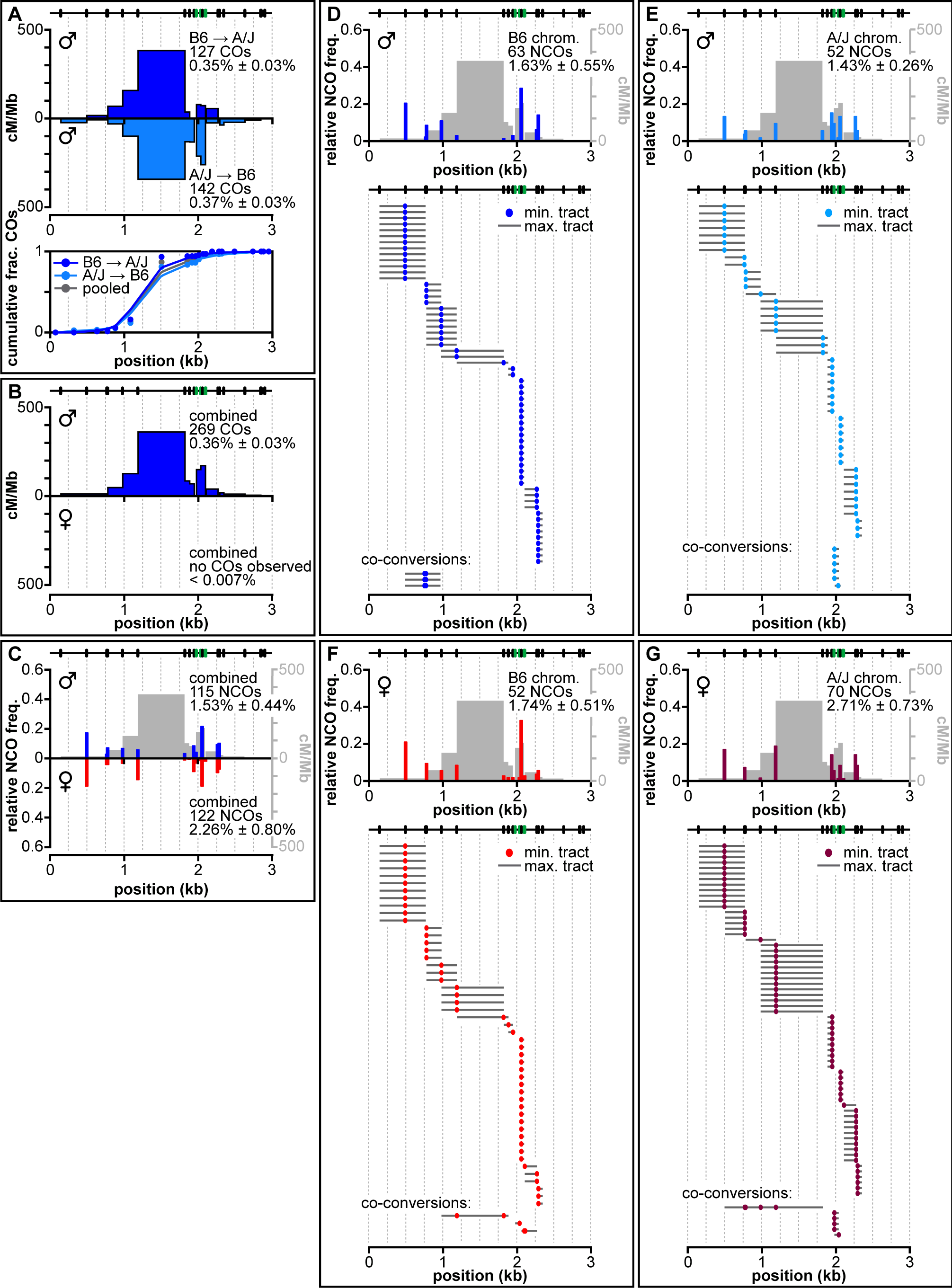
Recombination at the distal hotspot in males and females. **A.** Similar distributions of crossover breakpoints in males in both orientations. Crossover breakpoint maps and cumulative crossover distributions are shown as in Figure 1C. **B.** Difference in crossover formation between males and females. Data from both orientations of the allele-specific PCR were pooled for males and females, presented as in Figure 1D. No crossover molecules were recovered from oocyte DNA samples. **C.** Total noncrossovers in males and females, presented as in Figure 1E. **D–G.** Relative noncrossover frequencies and noncrossover gene conversion tract distributions in males (D,E) and females (F,G) on the B6 (D,F) and A/J (E,G) chromosomes, presented as in Figure 1F.

#### Noncrossovers

We identified 115 noncrossovers from ∼8,000 haploid genome equivalents of sperm DNA, for a Poisson-corrected frequency of 1.53% (♂, **Figure 2C-E**; **Table 1; Table S2**). Average minimum and maximum conversion tracts were 2 bp and 294 bp, respectively, comparable to *A3* (Cole et al. 2010) and the central hotspot, and co-conversions were scarce (**Figure 2D, 2E**). Three noncrossovers were detected that hybridized to two adjacent ASOs of the opposing allele. These co-conversions involved the same two polymorphisms 11 bp apart, located on the left flank of the hotspot (maximal tract 483 bp; **Figure 2D**). Six non-crossovers are presumptive co-conversions because they hybridized to ASOs that each contained two SNPs (polymorphisms 3 and 6 bp apart, **Table S2**).

The same PCRs yielded 25 crossover molecules (Poisson-corrected frequency of 0.33%; **Table 1**), matching expectation from the crossover-specific assay. The observed per-meiosis ratio is 1 crossover to 9.3 noncrossovers ((4 × 1.53% noncrossovers) ÷ (2 × 0.33% crossovers)). This is comparable to the ratio at *A3* (Cole et al. 2010). However, the middle of the hotspot contains a 634 bp stretch without polymorphisms between B6 and A/J haplotypes (**Figure 2, Table S2**). Thus, it is possible that noncrossovers occur frequently in this interval but escape detection.

### DSB levels and crossover frequencies are poorly correlated

To compare local DSB activity with recombination outcomes, we analyzed published data generated by deep sequencing of ssDNA bound by the DMC1 strand exchange protein in testis extracts from B6 animals (Brick et al. 2012). In this ssDNA sequencing (SSDS) assay, reads mapping to the forward and reverse strands represent DSB resection tracts to a DSB hotspot’s left or right side, respectively. Thus, although precise DSB distributions cannot be gleaned from these data, hotspot midpoints can be inferred to lie between the forward and reverse strand accumulations. Furthermore, total SSDS read count at a hotspot is expected to be proportional to DSB frequency (Pratto et al. 2014).

As expected, each hotspot displayed a cluster of SSDS reads centered on the midpoint of crossover activity (**Figure 3A**). However, the relative SSDS read count correlated poorly with the relative crossover frequency: compared to the distal hotspot, the central hotspot had less than one-third the frequency of SSDS reads but a 3.5-fold higher crossover frequency (**Figure 3A, Table 1**). The high DSB activity at the distal hotspot reinforces our suspicion that the actual noncrossover frequency at this hotspot is likely higher than we are able to detect due to the low polymorphism density in the hotspot center. More importantly, these findings imply that a higher DSB frequency does not necessarily translate into a higher crossover frequency.

**Figure 3.**
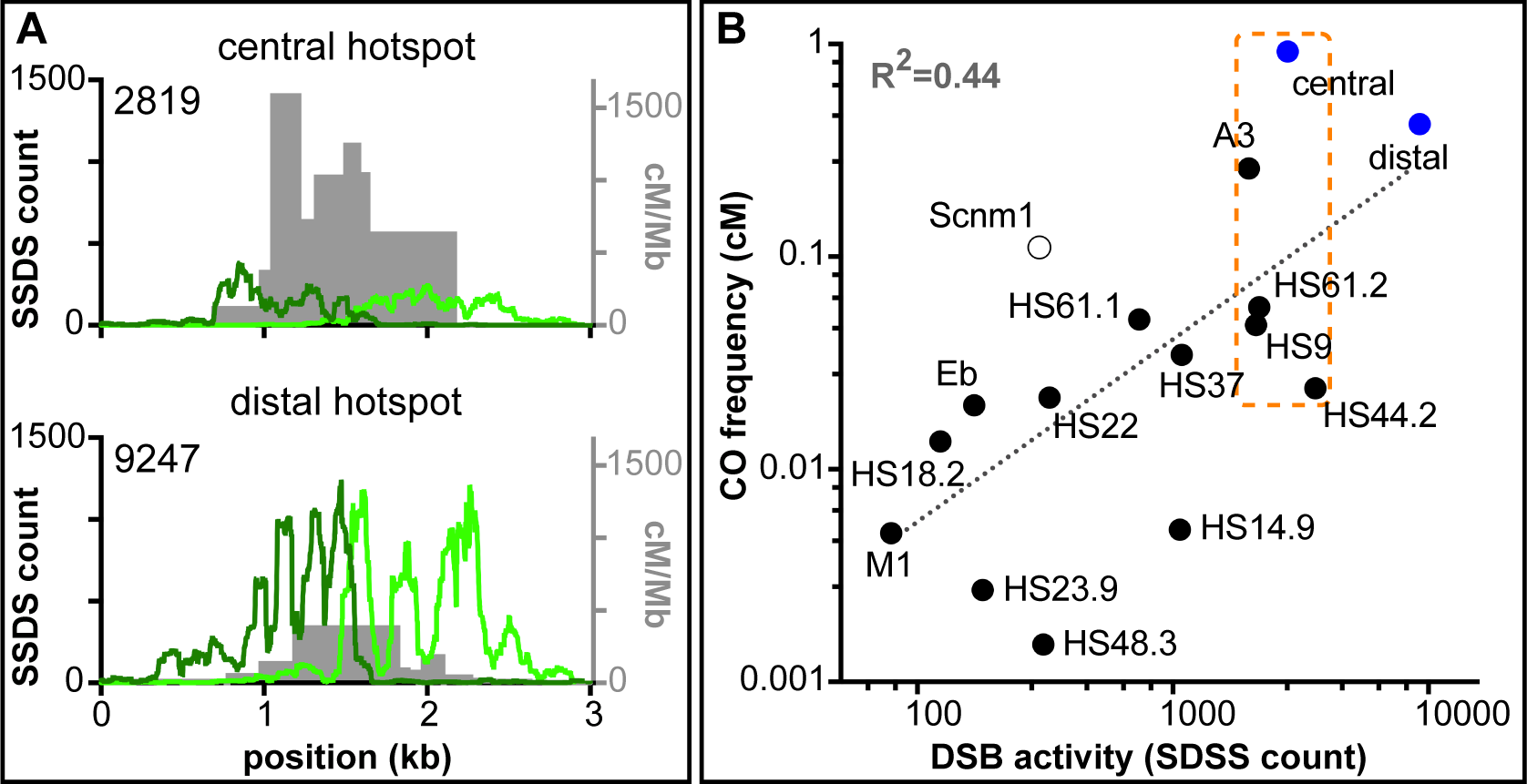
Estimates of relative DSB activity are an unreliable predictor of crossover frequency. **A.** Comparison of SSDS reads with crossover distributions at the central and distal hotspots. Crossover breakpoint maps from spermatocytes (pooled from both orientations of allele-specific PCR) are shown in gray. Forward-strand SSDS reads are shown in dark green and reverse-strand reads are in light green; total read counts (a measure of DSB activity) are indicated (data from Brick *et al*. (2012)). The midpoint between accumulations of the forward-and reverse-strand reads marks the hotspot center. Note that A/J and B6 share the same *Prdm9* allele, and the symmetry of crossover maps (Figures 1C and 2A) indicates that recombination initiation occurs at comparable frequencies on both haplotypes. Thus, the SSDS and crossover maps are directly comparable even though they were generated from animals of different strain backgrounds (pure B6 for SSDS vs. A/J × B6 F1 hybrids for crossovers). Note that the y-axis scales are the same for both hotspots. **B.** Comparison of SSDS read counts and crossover frequencies at published mouse hotspots. SSDS data are from Brick *et al.* (2012). Filled blue circles are the hotspots from this study; filled black circles denote published hotspots assayed by allele-specific PCR of sperm DNA; open circle denotes a published hotspot assayed for crossing over by pedigree analysis (see **Table S5** for details). Dotted line is a least-squares regression line fitted to the log-transformed data. The orange box marks a group of hotspots that are within a factor of two of each other for SSDS signal, but cover a much wider range in crossover activity. Note that different studies used different methods to correct recombination assays for amplifications efficiencies, so measured crossover frequencies may be underestimated to different degrees for specific hotspots (see Methods for further details).

To evaluate the generality of these findings, we compared SSDS read counts to published crossover frequencies determined by sperm typing or pedigree analysis at *A3* and other hotspots active in the B6 strain (**Figure 3B**). Although there was a positive correlation between SSDS counts and crossover frequency, the strength of the correlation was modest, with variation in the SSDS counts explaining less than half the variation in the crossover data (R^2^ = 0.44). Considering subsets of these hotspots is particularly revealing. For example, the hotspots highlighted in the box in **Figure 3B** are within a factor of two of each other for SSDS signal, but they cover a 38-fold range in crossover activity. A similarly poor correlation was seen for data for human hotspots, but emphasis was placed on existence of a correlation, not deviations of individual hotspots from the trend (Pratto et al. 2014). These findings strongly support the conclusion that the per-DSB crossover frequency can vary substantially between hotspots.

### The central hotspot is active in females, generating both crossovers and noncrossovers

To analyze recombination in females, we carried out the crossover and noncrossover/crossover assays on DNA extracted from ovaries of newborn B6 × A/J F1 hybrids. At this age, ovaries contain oocytes that are mainly in the late prophase stages of diplonema and/or dictyate arrest (Dietrich and Mulder 1983; McClellan et al. 2003). Analysis of the kinetics of interhomolog recombination in male mouse meiosis found that most crossovers and noncrossovers were formed by late pachynema, with little to no difference in timing between the two products (Guillon et al. 2005). Assuming comparable kinetics for female meiosis, oocytes from ovaries of newborns are expected to have largely completed meiotic recombination (Guillon et al. 2005; Baudat and de Massy 2007a). Thus, although we analyzed recombination in females at an earlier stage than in males, it is unlikely this difference would lead to underrepresentation of one or both recombination products.

Because most cells in ovary samples are somatic, we enriched for oocytes by disrupting the ovaries in the presence of collagenase and DNase I, followed by several wash steps to selectively lyse and deplete somatic cells (Eppig and Schroeder 1989; de Boer et al. 2013). We determined the fraction of oocytes in the enriched cell suspensions using immunocytology and corrected recombination frequencies accordingly (Baudat and de Massy 2007a; Baudat and de Massy 2009; de Boer et al. 2013).

#### Crossovers

: Allele-specific PCR in both orientations yielded 74 crossovers from a total input of ∼29,000 haploid genome equivalents from oocytes, for a Poisson-corrected overall crossover frequency of 0.23% (**Table 1**). No crossovers were detected in somatic controls (frequency < 0.002%). As in males, crossover breakpoints showed similar distributions for both orientations of allele-specific primers (**Figure 1H**), indicating no preference for either haplotype in recombination initiation (Jeffreys and Neumann 2002; Cole et al. 2010). Crossover breakpoints showed a similar distribution as in males (**Figure 1D**). Average activity was 244 cM/Mb, with a maximum of 1130 cM/Mb. The lower peak activity and overall frequency in females than in males may reflect a difference in crossover activity between the sexes. Alternatively, the observed difference could reflect different levels of precision in estimating absolute frequencies of recombinant DNA molecules from ovary vs. sperm samples. Nonetheless, this hotspot is highly active in females with an activity >400-fold greater than the genome average of 0.55 cM/Mb.

#### Noncrossovers

: From PCR in the universal-to-B6 orientation we recovered 14 noncrossovers from an input of 584 haploid genome equivalents from oocytes, for a Poisson-corrected frequency of 2.4% (**Figure 1I**; **Table 1; Table S1**), a frequency comparable to that of males. The average minimal and maximal conversion tracts were also similar to males (1 bp and 304 bp, respectively). No co-conversions were found among the 14 noncrossovers, agreeing with the low number in males. The majority of noncrossovers were within the central 2 kb of the hotspot, similar to that observed in males. We also retrieved two crossover molecules from the same assay (0.34%), consistent with the crossover assay. The observed per-meiosis ratio is higher than in males (1 crossover to 14 noncrossovers), but this difference should be viewed with caution as the small number of recombinants analyzed in females renders this estimate less precise. Overall, however, we can conclude that this hotspot behaves similarly in males and females.

### Crossovers are not detected at the distal hotspot in females

#### Crossovers

: In stark contrast to males, females displayed no detectable crossover activity at the distal hotspot. No crossovers were recovered from a total input of ∼15,000 haploid genome equivalents from oocytes by allele-specific PCR, for an overall crossover frequency of < 0.007% (♀, **Figure 2B; Table 1)**. Crossovers were also not detected in somatic controls (frequency < 0.004%). Because we could detect crossovers at the central hotspot in females, absence of crossovers at the distal hotspot cannot be ascribed to technical difficulty in detecting recombinants in oocytes, but must instead reflect a sex-specific difference in the behavior of this hotspot.

#### Noncrossovers

: To determine whether absence of crossovers reflects absence of recombination and thus likely DSBs, we analyzed noncrossover formation in oocytes using all four PCR orientations. From a total input of ∼6,000 haploid genome equivalents, we recovered 122 noncrossovers for a Poisson-corrected overall frequency of 2.3% (♀, **Figure 2C**; **Table 1; Table S2**). This value is comparable to the total recombination frequency in males, and both the spatial distribution and average conversion tract lengths of noncrossovers (minimal: 10 bp; maximal 345 bp) were similar in both sexes (♀ **Figure 2C, 2F, 2G**). Again, co-conversions were rare: three unambiguously detected co-conversions included more than one polymorphism (Figure 2F, 2G); seven noncrossovers involved presumptive conversion of more than one polymorphism contained within a single central ASO (**Table S2**). Most co-conversions were located at the right flank of the hotspot and were comparable to those found in males in that they involved two closely spaced polymorphisms (minimum conversion tract length of 5 bp, maximum of 208 bp). Two co-conversions had substantially longer conversion tracts than any events observed in males. One involved four polymorphisms, spanning at least 420 bp (maximum tract length 1324 bp); another involved two polymorphisms spanning 634 bp in the center of the hotspot (maximum tract length 903 bp). As in the crossover-specific assay, no crossovers were recovered from these PCRs (crossover frequency < 0.017%) (**Table 1**).

Thus, this hotspot is highly active for recombination initiation in female meiosis, despite the lack of detectable crossovers. These data strongly indicate that the sex-specific differences in crossover activity principally reflect differences in the recombination outcome between males and females rather than a difference in DSB number per se.

### Biased gene conversion in noncrossovers

The large number of noncrossovers identified at the distal hotspot in both males and females provided an opportunity to directly examine gene conversion bias at individual polymorphisms. Gene conversion is biased in favor of transmission of GC alleles in many eukaryotes (Duret and Galtier 2009). Such a bias has been observed directly in noncrossover (Odenthal-Hesse et al. 2014) and crossover (Arbeithuber et al. 2015) recombination products in human sperm, and can explain patterns of GC content enrichment at mouse hotspots (Clement and Arndt 2013). Accordingly, we found that in males noncrossovers frequently showed significant bias resulting in more conversion to GC than to AT (**Table S2**). Of the 8 GC/AT polymorphisms, 6 showed bias toward GC conversion. Normalizing for the number of chromosomes analyzed for noncrossovers, overall there were 64 conversions to GC but only 33 conversions to AT (p=0.0022, binomial test). Bias was observed throughout the hotspot, including at polymorphisms located 1 kb from the hotspot center. The more limited analysis at the central hotspot also showed a skewing toward GC conversion (29 conversions to GC, 15 conversions to AT, p=0.049, binominal test; **Table S1**).

In females, gene conversion bias was not as clear-cut, consistent with the suggestion that biased gene conversion is more prominent in males in humans than in females (Duret and Galtier 2009). Four of the 6 GC/AT polymorphisms that showed bias in males were also skewed toward GC conversion in females (**Table S2**). However, overall the bias was not significant (p=0.4926, binomial test; 56 conversions to GC and only 48 conversions to AT).

## DISCUSSION

Because of the uncertainties in precisely determining noncrossover frequencies, it has been difficult to know whether previously observed differences in crossover:noncrossover ratios between different hotspots (Holloway et al. 2006; Baudat and de Massy 2007b; Cole et al. 2010) reflect genuine differences in the crossover vs. noncrossover decision or are merely a result of shortcomings in noncrossover detection. In this report, we provide strong evidence for substantial variation in mouse interhomolog recombination outcomes (crossover vs. noncrossover). This lack of uniformity has important implications for understanding crossover control in complex genomes.

### Intrinsic differences between hotspots in the likelihood that a DSB will give rise to a crossover

In females, noncrossovers were detected at high frequency at both the central and distal hotspots, thus indicating substantial DSB formation at both hotspots, but crossovers, which can unambiguously be determined, were observed only in the central hotspot. Thus, irrespective of the exact frequency of noncrossovers, the central hotspot in females is markedly more biased towards crossover formation than the distal hotspot. In males, we find an analogous difference between the two hotspots by comparing relative crossover frequencies with inferred DSB levels: a 3.5-fold higher crossover frequency is observed at the central hotspot yet it has one-third the number of SSDS reads.

Analysis of published data for several hotspots located across the genome also shows that SSDS reads as a measure of relative DSB levels are a poor predictors of relative crossover frequencies, further supporting the conclusion that there are intrinsic differences between hotspots in the likelihood that a DSB will give rise to a crossover. A similarly modest correlation is obtained with an alternative method of estimating relative DSB levels, SPO11-oligonucleotide sequencing, suggesting that this pattern is not simply due to uncertainty in estimating relative DSB frequency from SSDS reads (R^2^ = 0.35; J. Lange, M. Jasin, and S. Keeney, unpublished observations).

In principle, the strong bias against interhomolog crossovers at the distal hotspot in females could be due to factors acting over a large chromosomal domain. For example, subtelomeric and/or centromeric regions in many species tend to have lower levels of crossover formation, with suppression of DSBs, changes in the crossover:noncrossover ratio or increased sister-chromatid recombination as suggested causes (Drouaud et al. 2006; Chen et al. 2008; Mancera et al. 2008). However, the distal hotspot is located at some distance (∼10 Mb) from the telomere and both male and female crossover hotspots have been mapped even closer to the telomere on the same chromosome (Paigen et al. 2008), such that a large-scale telomere effect is unlikely to be the sole reason for bias against crossing over at the distal hotspot. Instead, we speculate that factors operating more locally may play an important role. For example, whereas the central hotspot is intergenic, the distal hotspot is located within the *Rab3gap2* gene, with the center of the hotspot overlapping an exon (**Tables S1 and S2**).

### Modulation of the crossover:noncrossover decision in mouse

In males, the crossover:noncrossover ratios are ∼1:4 at the central hotspot vs. ∼1:9 at the distal hotspot. The short gene conversion tracts for most noncrossovers makes their detection highly dependent on the distribution of polymorphisms relative to DSBs (Cole et al. 2010). For the distal hotspot, the longest stretch without any polymorphisms (634 bp) is located at the center of the hotspot, which is where the peaks of crossover and noncrossover activity for most hotspots tend to overlap (Baudat and de Massy 2007b; Cole et al. 2010). In contrast, polymorphisms in the central hotspot are distributed more evenly, including at the hotspot center, where noncrossover activity is highest. Thus, although it is likely that noncrossovers are underestimated at both hotspots, we infer that the noncrossover frequency is underestimated to a greater degree at the distal hotspot. Even so, the strong bias towards noncrossover formation at this hotspot in males is much less extreme than that in females. While it remains possible that some DSBs in both hotspots are repaired by sister chromatid recombination as is known to occur in yeast (Schwacha and Kleckner 1994; Goldfarb and Lichten 2010; Hyppa and Smith 2010), these results provide clear evidence for crossover control at the level of the crossover:noncrossover decision.

Regional differences in recombination outcomes have been observed in the budding yeast *Saccharomyces cerevisiae* by whole-genome analysis (Mancera et al. 2008), but this study did not control for the density of sequence polymorphisms in hotspots, so differences in noncrossover detection may underlie some of the observed variation in recombination outcome (S. Keeney, unpublished observations). A separate study found that crossovers but not noncrossovers were substantially lower within 20 kb of chromosome ends compared with genome average (Chen et al. 2008), also suggestive of regional variation in recombination outcome, with recombination near telomeres relatively biased toward noncrossovers (Chen et al. 2008). However, non-allelic homologous recombination between dispersed repetitive sequences is known to exchange ends of different chromosomes at appreciable frequencies (Louis and Haber 1990; Louis et al. 1994), so these subtelomeric regions may be highly variable in yeast. If so, at least some of the observed reduction in crossover frequency may reflect large-scale structural differences between the ends of homologous chromosomes in the hybrid yeast strains analyzed. More recently, Borde and colleagues directly demonstrated examples of yeast hotspots with different DSB:crossover ratios (Serrentino et al. 2013). Greater propensity to form crossovers is correlated with enrichment for binding of the SUMO E3 ligase Zip3 (Serrentino et al. 2013). Although the factors that determine whether Zip3 will be enriched remain unknown, it is interesting to consider that tendency for enrichment of the Zip3 homolog RNF212 (Reynolds et al. 2013) might similarly affect recombination outcome in mouse.

The fission yeast *Schizosaccharomyces pombe* provides a distinct example for how recombination outcome can vary dramatically from place to place in the genome (Hyppa and Smith 2010; Fowler et al. 2014). In this organism, crossovers are distributed fairly uniformly, i.e., with nearly constant cM/kb, despite highly non-uniform distribution of DSBs (“crossover invariance”). DSBs in hotspots tend to be repaired more often using the sister chromatid as a template, so they are less likely to give rise to interhomolog crossovers or noncrossovers. In contrast, widely dispersed DSBs that form outside of detectable hotspots are usually repaired using the homolog as a template, and account for a disproportionately large fraction of crossovers. (An analogous bias toward using the sister chromatid may occur for recombination occurring near centromeres in budding yeast (Chen et al. 2008).) Available data have been interpreted to indicate that the crossover vs. noncrossover ratio for interhomolog events varies little between loci in fission yeast (Cromie et al. 2005).

Because we recovered numerous noncrossovers at the distal hotspot in both males and females, it is clear that interhomolog interactions via recombination are frequent here, implying that the biases in recombination outcome predominantly reflect variation in the choice of crossover vs. noncrossover pathways, not variation in the choice of homolog vs. sister chromatid as the template for repair. Thus, the regional variation in recombination partner choice that underlies crossover invariance in fission yeast is distinct from the variation in crossover vs. noncrossover outcome we observe in mice.

Our findings suggest parallels with studies of recombination outcomes at a human hotspot in the *PAR2* region of the X and Y chromosomes by allele-specific PCR in sperm DNA, where striking differences in crossover:noncrossover ratios were observed between men with similar or identical haplotypes within the hotspot (Sarbajna et al. 2012). This variation at a single hotspot seen when comparing genetically distinct individuals indicates that the choice of recombination outcome can be modulated by factors acting in trans or in cis but at a distance (outside the hotspot proper) (Sarbajna et al. 2012). We note that *PAR2* may not be fully representative of behavior of autosomal chromosome segments: it is a small region of shared homology at the distal tips of the short arms of the X and Y chromosomes. Unlike an autosomal segment, it is not flanked by large swathes of DNA capable of pairing and recombining with the homolog, but it is also unlike the longer *PAR1* region at the other end of the sex chromosomes, where crossing over occurs in nearly every meiosis and is critical for accurate sex chromosome segregation (Rouyer et al. 1986). Nonetheless, the behavior of PAR2 provides a clear example where hotspot context (in this case, genetic and/or epigenetic differences outside the hotspot) influences the likelihood that a DSB will give rise to a crossover. In broad strokes, this appears analogous to the sex-specific difference we observe for the distal hotspot.

We were intrigued that two of the co-conversions at the distal hotspot in females had unusually long conversion tracts that spanned or flanked the region where crossover activity in males was highest. Such long tracts have not been observed in males (Guillon et al. 2005; Svetlanov et al. 2008; Cole et al. 2010). Although the number of recovered events is small, the possibility arises that these co-conversions derive from a qualitatively different recombination intermediate than most other noncrossovers. For example, these events could be explained if they started out as double-Holliday junctions in the crossover pathway, but then became noncrossovers either by an unusual configuration of Holliday junction resolution or by Holliday junction dissolution. If so, this would imply the existence of an additional crossover control point at the double Holliday-junction stage, as has been proposed in yeast (Martini et al. 2011).

### Sex-specific crossover activity at the distal hotspot

In addition to differences in the crossover:noncrossover ratio between hotspots, our study also demonstrates an example of an extreme difference in behavior of a single hotspot when assayed in the different cellular contexts of the oocyte and spermatocyte. Based on both the frequency and distribution of noncrossovers, the distal hotspot can be considered as comparably highly active in females and males, yet crossovers were observed only in males. To our knowledge, this is the first direct evidence of intrinsic bias in recombination outcome in mammalian meiosis, and the first evidence in any organism that such bias can change in a cell-type specific manner. It is interesting in this regard that males have a higher likelihood of undergoing crossovers in the centromere-distal regions of chromosomes compared with females. Other crossover-suppressed hotspots may be uncovered in the distal portion of chromosomes in females. Mouse pedigree studies mapping crossovers in both male and female meioses on chromosomes 1 and 11 identified numerous hotspots. Some hotspots were sex-specific, and some of the hotspots shared between the sexes showed different crossover frequencies in males vs. females (Paigen et al. 2008; Billings et al. 2010). So, while the overall propensity towards crossing over is greater in males in the centromere-distal region, the crossover to noncrossover ratio will likely vary considerably between individual hotspots.

Our finding of intrinsic differences in crossover vs. noncrossover frequencies, both between hotspots and at the same hotspot between males and females, provides new insight into the degree to which recombination outcome can be locally regulated in mammals. Although the presence of a DSB is an absolute prerequisite for crossover formation, our findings show that high DSB levels do not guarantee a high crossover activity or, as exemplified by the distal hotspot in females, any crossover activity at all. Thus, hotspot activity is regulated at multiple points from DSB formation through to the crossover/noncrossover decision.

## MATERIALS & METHODS

### Mouse strains

The A/J x C57BL/6J F1 hybrids used in this study were either directly purchased (males) or bred from strains from the Jackson Laboratory (females and males). All experiments were done according to relevant regulatory standards and were approved by the MSKCC Institutional Animal Care and Use Committee.

### DNA extraction

DNA extractions were performed as described (Kauppi et al. 2009; Cole and Jasin 2011; de Boer et al. 2013). DNA was extracted and analyzed from 2 males and 2 pools of ∼45 females. Sperm DNA was extracted from cauda epididymides from adults. Ovary DNA was extracted from newborns, which were born on day 19–21 of gestation. A cell suspension was made from collected ovaries and enriched for oocytes (Eppig and Schroeder 1989; de Boer et al. 2013). A small aliquot of ovary cell suspension was used for immunocytological labeling with anti-SYCP3 to determine the fraction of oocytes (Baudat and de Massy 2007a; Baudat and de Massy 2009; de Boer et al. 2009), which averaged 35%. Liver DNA from the same mice that provided the sperm or ovary DNA served as a negative control.

### Hotspot identification and confirmation

The central hotspot was identified using A/J x C57BL/6J recombinant inbred strains as described (Bois 2007) (See also **Supplemental Information**). Information about the distal hotspot was generously provided by G. Petukhova and R.D. Camerini-Otero (Smagulova et al. 2011). SNPs from Shifman *et al*. (Shifman et al. 2006) and the dbSNP database (NCBI) were confirmed and additional SNPs and indels were identified by sequencing genomic DNA from both parental strains (Jackson Laboratory; primers in **Tables S3A and S4A**). Allele-specific and universal PCR primers for both hotspots were designed and optimized as described (Kauppi et al. 2009). (Universal primers in **Tables S3B and S4B;** allele-specific primers in **Tables S1 and S2**). Amplification efficiency for each DNA sample and each primer (allele-specific and universal) was determined by performing 16 PCRs with inputs of 12, 24 and 60 pg DNA per reaction, for each allele-specific PCR primer against a universal PCR primer (Cole and Jasin 2011). An initial test of hotspot activity was conductured by amplification of crossover molecules as described (Kauppi et al. 2009; Cole and Jasin 2011), with pools ranging from 300 to 3000 input DNA molecules. Liver DNA was used as a somatic (negative) control at equivalent total input DNA.

### Recombination assays

For the crossover-specific assay, amplification of recombinant molecules using allele-specific primers (**Figure 1Bi**; **Tables S1 and S2**) and pools of 150–400 input DNA molecules was performed as described (Kauppi et al. 2009; Cole and Jasin 2011). Crossover assays were performed in both orientations; liver DNA and no-input DNA were used as negative controls. Crossover frequencies were corrected for amplification efficiency and calculated (with estimates of standard deviation) using Poisson correction as described (Baudat and de Massy 2009), except that amplification efficiency was considered a constant for each set of primers. Crossover-positive PCRs and negative controls were re-amplified using nested universal PCR primers (**Tables S3B and S4B**), PCR products were transferred onto nylon membranes, and crossover breakpoints were mapped by hybridization with allele-specific oligos (**Tables S1 and S2**) as described (Kauppi et al. 2009).

For the noncrossover/crossover assay, recombinant molecules (along with nonrecombinant molecules of the selected haplotype) were amplified using allele-specific PCR primers against universal PCR primers with pools of 15 input DNA molecules (**Figure 1Bii**; **Tables S1, S2, S3B and S4B**), as described (Kauppi et al. 2009; Cole and Jasin 2011). Assays were performed in all four possible primer orientations, except for the central hotspot in females.

PCR products were transferred onto nylon membranes, and noncrossovers and crossovers were detected and mapped by hybridization with allele-specific oligos as described (Kauppi et al. 2009; Cole and Jasin 2011). Noncrossover and crossover frequencies were corrected for amplification efficiency and calculated using Poisson correction as described for the crossover-assay. For both calculation of the overall noncrossover frequencies and the graphical representations of noncrossovers across the hotspots, co-conversions were normalized by dividing by the number of polymorphisms involved.

For females, the observed crossover and noncrossover frequencies were corrected for the fraction of oocyte-derived amplifiable molecules for each ovary DNA sample as described (Baudat and de Massy 2009), except that the correction factor was considered a constant.

### Comparison of SSDS read counts and crossover frequencies at published mouse hotspots

SDSS data were from Brick *et al*. (2012). Crossover frequencies were from (Buchner et al. 2003; Yauk et al. 2003; Bois 2007; Kauppi et al. 2007; Cole et al. 2010; Wu et al. 2010) (see also **Table S5**). The crossover data used are for F1 hybrids involving the B6 strain background. With the exception of the central, distal and *A3* hotspots, correction factors for amplification efficiencies were not determined separately for specific primer pairs, so crossover frequencies may be underestimated to different (unknown) degrees in data from different sources. However, from modeling of the effects of additional correction factors comparable to those we observed in this study, it is unlikely that this uncertainty is a substantial contributor to the weakness of the regression relationship shown in **Figure 3B** (data not shown). For some hotspots, reciprocal crossover asymmetry has been observed, indicative of different DSB frequencies on the two haplotypes in the F1 hybrid assayed (**Table S5**). For example, HS22 displayed reciprocal crossover asymmetry in the B6 × DBA F1 hybrid, with the orientation of asymmetry indicating that DSB formation is more frequent on the B6 chromosome (Bois 2007). This implies in turn that relative DSB activity in the F1 hybrid is overestimated by using SSDS data from a pure B6 background. Importantly, correcting for this would not improve the overall regression relationship but would instead make it worse, so the overall poor relationship between SSDS frequency and crossover frequency is not simply a consequence of comparing data derived from different strain backgrounds.

## Acknowledgements

We are grateful to Galina Petukhova and Julian Lange for sharing data prior to publication, and to Liisa Kauppi and Francesca Cole for advice on performing allele-specific PCR assays for meiotic recombination. This work was supported by NIH grant R01 HD053855 (to S.K. and M.J.). E.d.B. was supported in part by Netherlands Organization for Scientific Research Rubicon Grant 825.07.006. S.K. is an Investigator of the Howard Hughes Medical Institute.

## SUPPLEMENTAL FIGURE LEGEND

**Figure S1.**
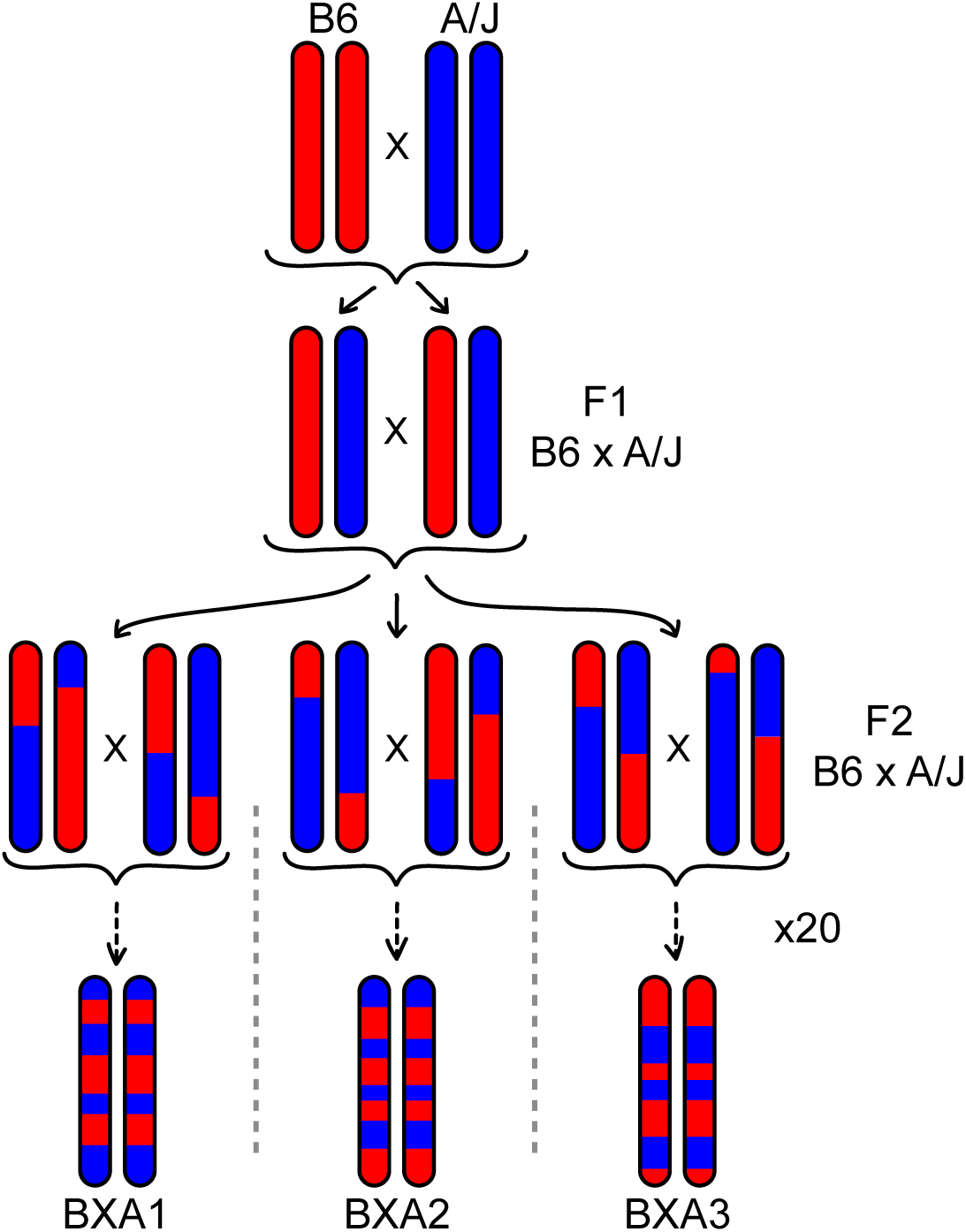
Breeding scheme for generating recombinant-inbred strains. RI strains are named using abbreviations for the maternal and paternal progenitor strains. Each RI strain can be viewed as a library of crossovers that occurred during the generations of inbreeding prior to fixation of homozygosity. Adapted from Bois (2007).

